# Accurate *de novo* design of a voltage-gated anion channel

**DOI:** 10.1101/2024.12.25.630309

**Authors:** Chen Zhou, Huican Li, Jiaxing Wang, Zhilin Chu, Shijin Sun, Ke Sun, Aiqin Zhu, Jiawei Wang, Xueqin Jin, Fan Yang, Tamer M. Gamal El-Din, Kun Wu, Peilong Lu

## Abstract

Ion channels gated by environmental cues play key roles in fundamental biological processes. Designing ion channels with tailored gating mechanisms remains a significant challenge due to the complexities involved in designing conformational changes in proteins. Here we report the accurate *de novo* design of a voltage-gated anion channel, namely VGAC. VGAC adopts a 15-helix pentameric structure featuring a constriction composed of five arginine residues within the transmembrane span. In patch-clamp experiments, VGAC showed strictly voltage-dependent currents and demonstrated selectivity for chloride anions over iodide anions. Our data suggests that the arginine constriction undergoes voltage-induced conformational changes, serving as both the voltage sensor and selectivity filter. A 2.9-Å-resolution cryo-electron microscopy structure of VGAC closely aligns with the design model. The ability to design ion channels with custom-designed conformational changes provides new insights into our understanding of the fundamental principles of membrane biophysics and unveils a wide range of potential applications.

## INTRODUCTION

Transmembrane proteins facilitate the exchange of ions, metabolites, and thus information across the cell membrane, thereby enabling cells to adapt more effectively to the surrounding environment. Although progress has been made in designing a Zn^2+^/H^+^ antiporter^1^, stable transmembrane protein scaffolds^2–4^, and transmembrane pores^5–9^, the *de novo* design of transmembrane proteins that can specifically respond to stimuli through induced conformational changes remains a significant challenge^8,10–14^. Specifically, *de novo* design of voltage-gated ion channels, which regulate the passage of specific ions across the membrane via conformational changes responding to changes in membrane potential, remains a formidable obstacle yet to be overcome.

Voltage-gated ion channels enable cells to shape electrical signals, which are crucial for the function of excitable cells such as neurons, muscles, pancreatic beta cells, and select types of immune cells^15^. Voltage-gated sodium, potassium and calcium channels share a similar overall architecture, consisting of four voltage-sensing domains (VSDs) that encompass a central ion conduction pathway^16–18^. The fourth helix (S4) of the VSD contains an array of positively charged residues, which serve as the primary structural elements responsible for voltage sensing. Changes in voltage potential across the membrane induce the movement of the S4 helix relative to the rest of the protein, leading to conformational changes that facilitate channel gating^19^. Voltage-gated chloride channels (CLC channels) possess an architecture that differs from the classic architecture of voltage-gated cation channels, instead resembling Cl^−^/H^+^ transporters^20^. The fast-gating response of CLC channels to voltage involves the conformational change of the gating glutamate side chain^21^.

To design voltage-gated ion channels, three interconnected design challenges must be addressed. Firstly, it is crucial to incorporate charged residues into the transmembrane spans for voltage sensing, a concept derived from naturally existing voltage-gated ion channels. However, this incorporation introduces a significant solvation energy cost when these charged residues are embedded in the membrane or within the protein core. These charged residues must either be balanced by countercharge residues or made solvent accessible. Secondly, it is imperative to consider the conformational changes that take place in the charged gating residues, as well as the interaction between the gating charges and other regions of the protein when exposed to different membrane potentials. Lastly, it is necessary to ensure that the core residues are well packed to stabilize the channel structure and allow for a water-accessible pore in the membrane while minimizing non-specific hydrophobic interactions with lipids or lipid-facing residues.

Here, we report the accurate *de novo* design of a voltage-gated anion channel, namely VGAC. VGAC was designed to feature a pentameric funnel-like structure, wherein each protomer consists of three transmembrane helices. A positively charged constriction composed of five arginine residues was designed within the transmembrane span, serving as both the voltage sensor and selectivity filter for anions. Moreover, VGAC adopts an Nin-C_out_ topology on the membrane, with the N- and C-termini located inside and outside the cell, respectively. VGAC exhibits strong selectivity for chloride ions and demonstrates voltage-dependent currents with strong outward rectification in whole-cell patch-clamp experiments. Single-channel recordings showed that the opening of the VGAC channel is regulated by membrane potential. The open probability dramatically increases at membrane potentials above +40 mV, likely due to the voltage-induced opening in the arginine constriction. The open-channel conductance of VGAC was determined to be 41.98±2.71 pS, which is comparable to some naturally existing ligand-gated chloride channels^22^. A 2.9-Å-resolution cryo-electron microscopy (EM) structure of VGAC closely aligns with the design model. Our findings pave the way for the development of a new generation of transmembrane proteins capable of responding to specific environmental cues through custom-designed conformational changes.

## RESULTS

### De novo design of funnel-shaped 15-helix pentameric structures

We focused on designing 15-helix transmembrane channels that display five-fold symmetry and form two concentric rings, referred to as TMH3C5 (Figure S1A). Each subunit consists of three helices connected by two short loops. The second helix, designated as S2, constitutes the inner ring, whereas the other two helices (S1 and S3) contribute to the outer ring. In comparison to helical transmembrane pores and channels that have been previously designed from scratch^5–8^, wherein the repeating subunit is predominantly composed of one or two helices, the TMH3C5 scaffold displays higher structural complexity and a greater density of interhelical interactions (Figure S1A), which may favor the incorporation of charged residues and accommodating conformational changes.

We first designed 15-residue helical motifs responsible for the formation of the inner ring of the channel (Figures 1A and 1B). We extended the Crick coiled-coil parameterization method^23–25^ to sample α-helical and super-helical parameters. In addition, we also sampled the tilt angle of the α-helix around y-axis. This method enables us to sample a greater variety of structural space, capturing not only cylindrical helical bundles but also funnel-shaped ones (see STAR Methods). We employed ProteinMPNN^26^ for sequence design and subjected the designed sequences to AlphaFold2^27^ (AF2) predictions, allowing us to evaluate the extent to which these sequences encoded the desired ring structures. From the pool of 1,207 sequences, we identified 8 sequences for which the predicted structures closely resembled the target designs. Notably, these structures clustered into two distinct geometries: a cylinder and a funnel (Figures 1B, S1B and S1C).

**Figure 1.**
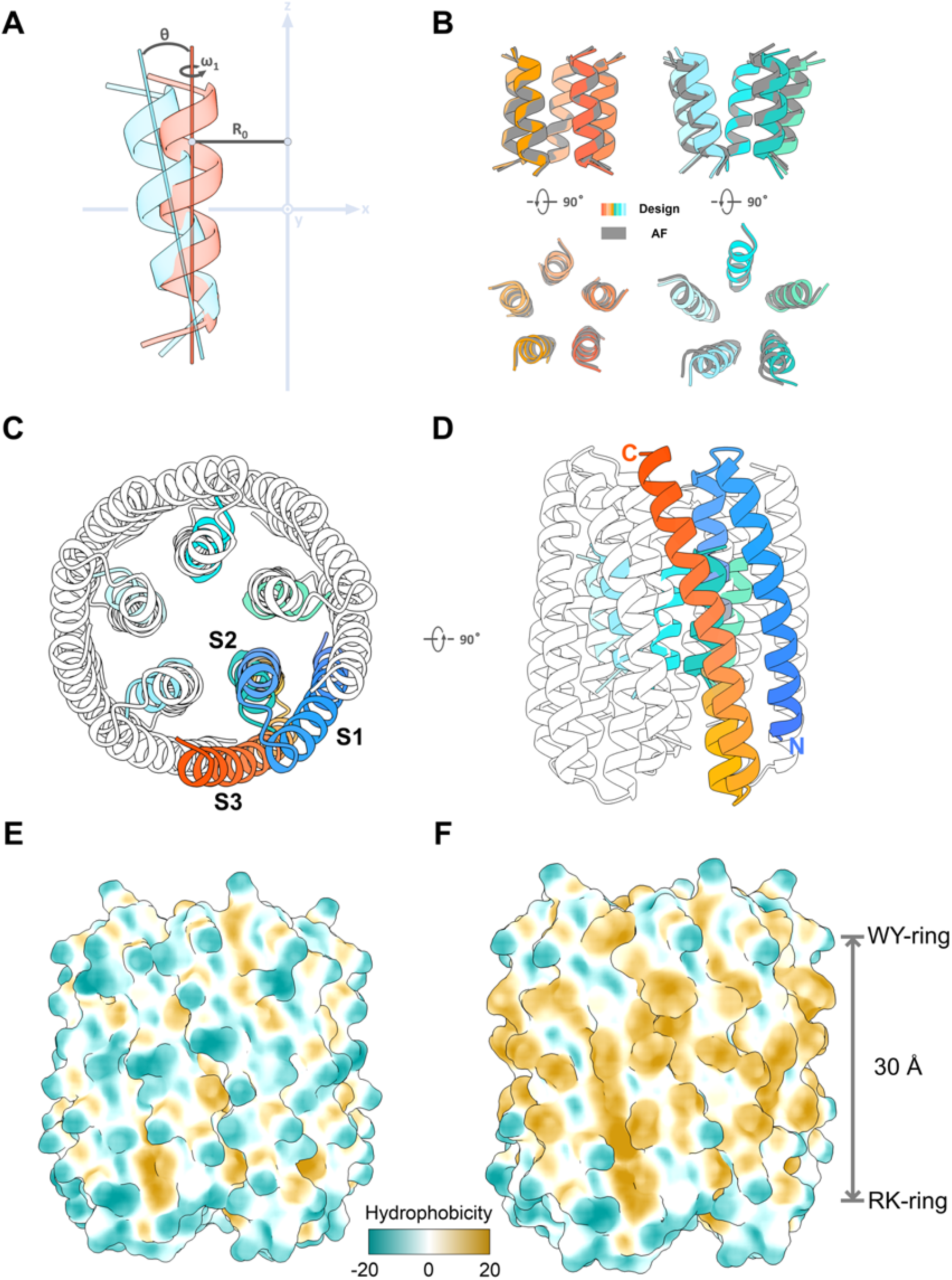
Design of 15-helix pentameric transmembrane channels. **(A)** Geometric interpretations of the sampled parameters for the inner-ring helical bundle. Geometrical meanings of the superhelical radius (R0) and starting helical phase (Δω1) in the Crick coiled-coil parameterization and the tilt angle (θ) of the α-helix around the y-axis are shown. The diagram illustrates a Cartesian coordinate system, where the x-z plane is depicted as the plane of the paper, and the y-axis emerges vertically from it. The z-axis represents the central C5 symmetry axis. The red and blue lines in the diagram represent the local helical axis. When the θ value equals 0, a cylinder helical bundle with a fixed superhelical radius is generated. Deviation of the θ value from 0 indicates a rotation of the local helical axis around the y-axis, resulting in the formation of a funnel-like shape, characterized by varying superhelical radii along the central axis. **(B)** Representative examples of the designer helical motifs strongly predicated to fold to the target structure by AlphaFold2 (AF2). These structures clustered into either a cylindrical or a funnel-shaped geometry at a TM-score clustering threshold of 0.6. **(C and D)** The top (**C**) and side (**D**) views of a representative designer WSH3C5 featuring two concentric rings. This helical-bundle backbone was built on a funnel-shaped motif (colored cyan) using the fragment assembly approach. One protomer is highlighted and colored spectrally from blue at its N-terminal segment to red at its C-terminal segment. **(E and F)** Scheme of the conversion from a WSH3C5 (**E**) into a TMH3C5 (**F**). The color of the surfaces corresponds to the degree of hydrophobicity, with green, white, and yellow denoting low, moderate, and high hydrophobicity, respectively. The transmembrane helical bundle contains a hydrophobic transmembrane span of approximately 30 Å. See also Figure S1.

Based on these two types of motifs, we constructed helical-bundle backbones using the fragment assembly approach^28^ (Figures 1C and 1D, S1D). This involved creating "blueprints" that specified the lengths of the helices and connecting loops for each scaffold. Ensemble of conformations for each blueprint was then generated through Monte Carlo-based assembly of short protein fragments. We employed ProteinMPNN^26^ to design the amino acid sequences for water-soluble helical bundles (WSH3C5, see STAR Methods). We conducted AF2 structure prediction calculations for each designed sequence and identified 27 designs that closely matched the predicted structures.

### De novo design of voltage-gated ion channels

Next, we designed the transmembrane helical bundles from the water-soluble design models (Figures 1E and 1F). To facilitate successful membrane localization, we utilized Rosetta to redesign the lipid-exposed residues, ensuring that the revised sequence possessed the essential characteristics for membrane localization, such as the hydrophobic transmembrane span, RK-ring, and WY-ring^2^. The design sequences are distinct from native protein sequences, as no matches were identified in a search against the non-redundant protein sequences database using BLAST^29^ (http://blast.ncbi.nlm.nih.gov/). Subsequently, synthetic genes were obtained for 8 transmembrane designs that closely matched the models predicted by AF2 (Figures S1E and S1F). These transmembrane proteins were then expressed in *E. coli* and purified from the membrane fraction. During size exclusion chromatography (SEC), two designed proteins, namely tmZC6 and tmZC8, can be purified to homogeneity (Figure S2A, S2B, S2D and S2E). The circular dichroism spectrum of tmZC6 and tmZC8 showed that the structures were helical and stable to thermal denaturation up to 95 °C (Figures S2C and S2F). To facilitate the determination of the cryo-EM structure of designer proteins, we fused the homo-pentameric BTB domain of KCTD1 protein^30^ to the N-terminus of the tmZC8 protein. This fusion construct was named tmZC8-BTB (Figure S1G). However, fusion to the tmZC6 design did not yield a qualified design that passed the AF2 prediction. The intervening linker between tmZC8 and BTB domain composed of five residues, which was designed as a short rigid helix using RosettaRemodel^28^. tmZC8-BTB exhibits a flower-like architecture with a height of approximately 100 Å, consisting of a transmembrane base and a corolla-shaped soluble domain. The purified tmZC8-BTB eluted as a single peak in SEC (Figures S2G and S2H).

Next, we sought to design the structural elements in tmZC8-BTB to achieve voltage sensitivity and ion specificity. Lysine (Lys) and arginine (Arg) residues carrying positive charges were introduced into the pore (Figure 2A), to serve two main functions. Firstly, these residues create a positively charged electrostatic environment within the pore, thereby attracting anions. Secondly, these residues could move uniformly in one direction in response to membrane potential, which may enable gating by adjusting the aperture of the funnel-shaped channel. The aperture of the channel is determined by several factors of the gating residues, including the charged state, size, rotamer state, relative position within the pore, and interaction with other parts of the pore under various membrane potentials. In our design, we propose these positively charged residues may serve as both the voltage sensor and selectivity filter for anions. One design, tmZC8-3R-BTB, incorporated three Arg residues at positions 157, 161, and 165 (Figures 2A and S3A). The purified tmZC8-3R-BTB eluted as a single peak in SEC, with an elution volume similar to that of tmZC8-BTB (Figures S2G-S2J).

**Figure 2.**
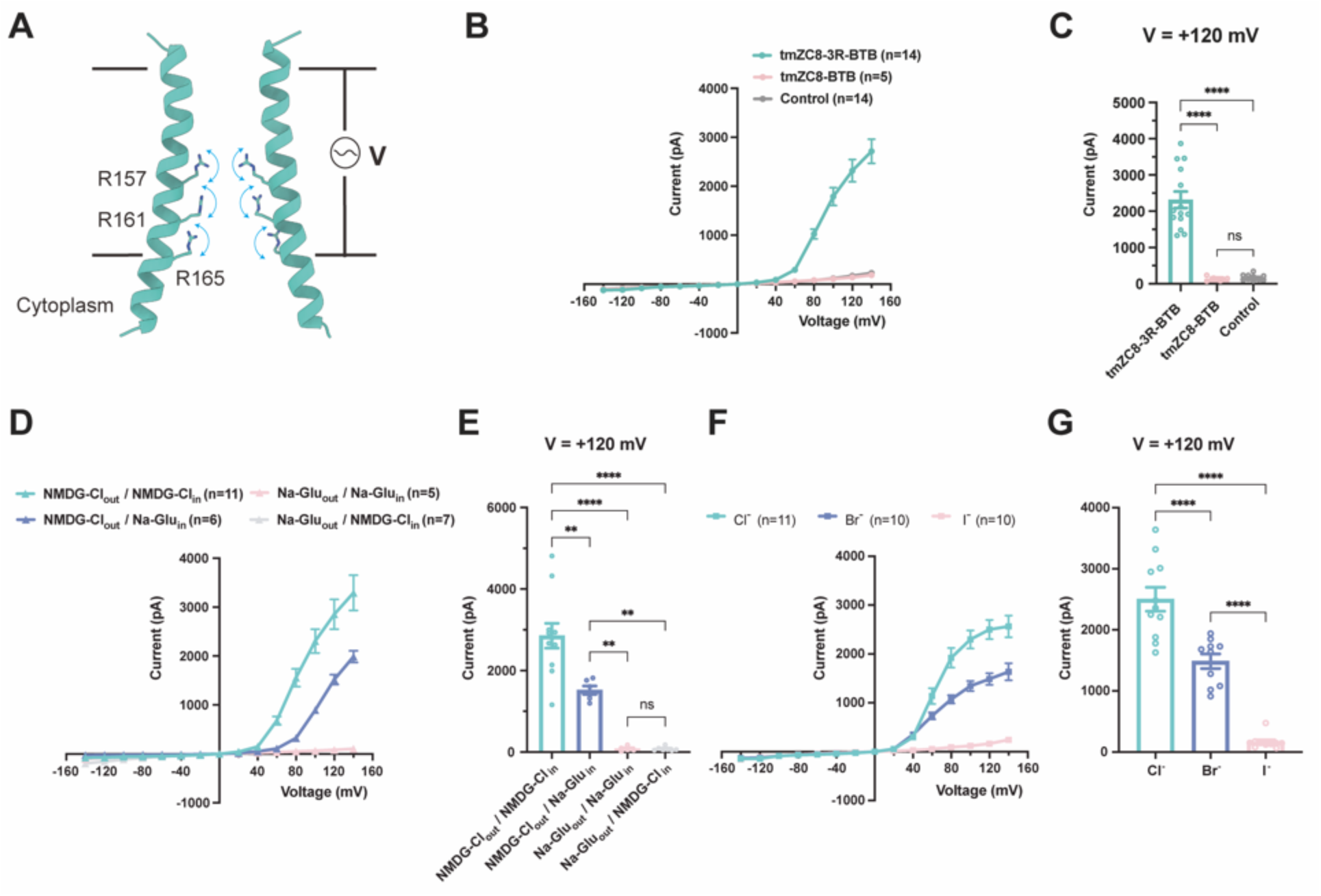
Design and characterization of a voltage-gated anion channel. **(A)** Design principles for voltage-gated ion channels. The charged side chains can assume distinct conformational states across varying membrane potentials. In a simplified scenario, positively charged residues can exist in a "down" state under negative potentials (negative inside) and an "up" state under positive potentials (positive inside). The movement of these charged residues in response to the membrane potential allows for gating by adjusting the aperture of the funnel-shaped channel. **(B and C)** Conductivity in whole-cell patch-clamp experiments in CHO-K1 cells expressing tmZC8-3R-BTB (green) or tmZC8-BTB (pink). The ion conductance of cells transfected with an empty vector is shown as a control (gray). **(B)** I-V curves showing a voltage-dependent increase in current starting from +40 mV. The number of measured cells is indicated in parentheses. **(C)** Bars represent the mean currents at 120 mV for two designs compared to control conditions. **(D and E)** Conductivity of tmZC8-3R-BTB with solutions containing Na-Glu or NMDG-Cl in the bath (out) and pipette (in) solutions. **(F and G)** Conductivity of tmZC8-3R-BTB with the bath (out) solutions containing different anions (chloride, Cl^-^; bromide, Br^-^; iodide, I^-^; all with sodium as the cation). The pipette (in) solutions contained NMDG-Cl. Error bars depict in **(C, E, G)** the mean ± standard error of the mean (SEM). The P-values obtained from the one-way ANOVA (Analysis of Variance) followed by Tukey’s comparison test are reported (*p < 0.05, **p <= 0.01, ***p <= 0.001, ****p <= 0.0001, ns = no significance). See also Figure S4 and Table S1.

### Nin-C_out_ topology on membrane

Both tmZC8-BTB and tmZC8-3R-BTB monomers were predicted by TMHMM^31^ to display a three-transmembrane-span topology, in which the N-terminal is located inside the cell and the C-terminal is outside the cell, consistent with the design model (Figures S3B and S3C). To examine the membrane localization and transmembrane topology of tmZC8-BTB and tmZC8-3R-BTB, fluorescence confocal imaging was conducted on *Xenopus* oocytes expressing designs tagged with C-terminal enhanced green fluorescent protein (EGFP, Figures S3D-S3F). The data shows that tmZC8-BTB and tmZC8-3R-BTB localized on the plasma membrane of the oocyte cells. The C-terminal EGFP can be stained by an anti-EGFP nanobody added outside the cell, suggesting that the C-termini of tmZC8-BTB and tmZC8-3R-BTB are located on the extracellular side of the membrane. Additionally, flow cytometric analysis was performed on human embryonic kidney 293T (HEK293T) cells expressing either tmZC8-BTB or tmZC8-3R-BTB with an EGFP tag at the C- or N-terminal (Figures S3G-S3J). The intracellular or extracellular localization of the EGFP tag in these cells was investigated by incubating the cells with a Cy5-labeled anti-EGFP nanobody. The results showed that cells expressing tmZC8-BTB or tmZC8-3R-BTB tagged with C-terminal EGFP can be stained by the anti-EGFP nanobody, but not the ones with the N-terminal EGFP. Taken together, these results show that tmZC8-BTB and tmZC8-3R-BTB adopt an Nin-C_out_ topology on the membrane.

### Whole-cell patch-clamp experiments showed time and voltage-dependent currents

Whole-cell patch-clamp experiments were performed on Chinese hamster ovary K1 (CHO-K1) cells expressing tmZC8-3R-BTB or tmZC8-BTB. Different ionic solutions were present in the extracellular (bath) and intracellular (electrode) solutions. In our experiments, tmZC8-3R-BTB showed time and voltage-dependent currents characterized by strong outward rectification (green line in Figures 2B and 2C; Figure S4A, S4D and S4E; STAR Methods). These currents remained within the background level in the voltage range of −140 mV to 40 mV. However, at potentials above 40 mV, the current rapidly increased, reaching 2319±227 pA at +120 mV. The kinetics of voltage-dependent activation of outward currents were faster when applying more depolarized potentials. This decrease in the activation time constant with more depolarized potentials gives an estimate of the speed of opening of the activation gate (Figure S4E). This behavior is similar to some naturally existing ion channels like K2P channels^32^ and the proton-activated chloride channels (PAC^33^). tmZC8-BTB did not show ionic current distinguishable from the background level in any of the different solutions used, indicating that tmZC8-BTB does not facilitate ion conduction (pink line in Figures 2B and 2C; Figure S4A-S4C; STAR Methods). To rule out any unexpected effect of the BTB domain, we further eliminated the soluble BTB domain from tmZC8- 3R-BTB, resulting in a construct namely tmZC8-3R. Subsequent whole-cell patch-clamp recordings revealed that tmZC8-3R exhibited time and voltage-dependent currents resembling of tmZC8-3R-BTB, indicating that the BTB domain has minimal impact on the observed currents (Figures S4F and S4G).

### Ion selectivity of the designer channel

We hypothesized that tmZC8-3R-BTB may function as an anion-selective channel due to the presence of arginine constriction. Thus, we conducted the patch clamp experiments using extracellular and intracellular solutions containing N-methyl-D-glucamine chloride (NMDG-Cl, NMDG^+^ cation with a radius of ∼3.65 Å^34^ which generally does not permeate through ion channels). The results showed strongly outwardly rectifying currents (green lines in Figures 2D and 2E; Figure S4H; Table S1), resembling that shown in Figure 2B (green lines). This indicates that the channel allows the influx of Cl^−^. Subsequently, we maintained the presence of NMDG-Cl in the bath solution while switching the pipette solution to sodium gluconate (Na-Glu, Glu^-^ anion with a diameter of ∼5.9 Å^35^ which generally does not permeate through ion channels) (blue lines in Figures 2D and 2E; Figure S4H; Table S1). This alteration resulted in a rightward shift in the I-V curve by approximately 40 mV, with mean currents of 1,518±102 pA at +120 mV. This may result from a partial block of the activation gate by gluconate anion that gets removed with very positive potentials. Strikingly, no current was detected with NMDG-Cl in the pipette solution and Na-Glu in the bath solution (gray lines in Figures 2D and 2E; Figure S4H; Table S1). When Na-Glu was present in both the bath and pipette solutions, no current was detected (pink lines in Figures 2D and 2E; Figure S4H; Table S1). These results suggest that the designer channel facilitates the influx of Cl^−^ into the cell while impeding the passage of Na^+^, NMDG^+^, and Glu^−^. These findings are consistent with our initial hypothesis, suggesting that the presence of positively charged Args in the pore enables the transport of anions. Interestingly, no current was observed at negative potential, which suggests that the Cl^−^ current is unidirectional. This indicates the closed gate of the designed channel at negative potential, blocking the efflux of chloride ions.

We evaluated the selectivity for anions (Cl^−^, Br^−^ and I^−^) (Figures 2F, 2G, and S4I; Table S1). The channel displayed significantly higher conductance for Cl^−^, with mean currents of 2501±197 pA at +120 mV. The ion selectivity order was Cl^−^ (2501±197 pA) > Br^−^ (1486±119 pA) ≫ I^−^ (163±36 pA) at +120 mV. Additionally, to investigate the potential impact of cation content in the pipette solution on the influx of chloride ions, experiments were conducted using NMDG-Cl, LiCl, NaCl, and CsCl solutions with the same protocol. Consistently, all four solution conditions exhibited similar currents, showing notable outward rectification. The observed Cl^−^ current is minimally affected by the cation content in the solution (Figures S4J and S4K; Table S1). Collectively, these results establish that tmZC8-3R-BTB functions as a voltage-gated anion channel, and it has been consequently renamed VGAC.

### Single-channel recordings of VGAC channel

To test the function of VGAC at the single-molecule level, we performed single-channel recordings of VGAC channels expressed in CHO-K1 cells using the patch-clamp technique in the outside-out configuration. Application of various transmembrane potentials (30 mV, 60 mV, 90 mV, and 120 mV) resulted in well-resolved unitary currents with a linear single-channel I-V relationship (Figures 3A and 3B), confirming that VGAC is indeed an ion channel. The single-channel conductance of VGAC was determined to be 41.98±2.71 pS (Figure 3B).

**Figure 3.**
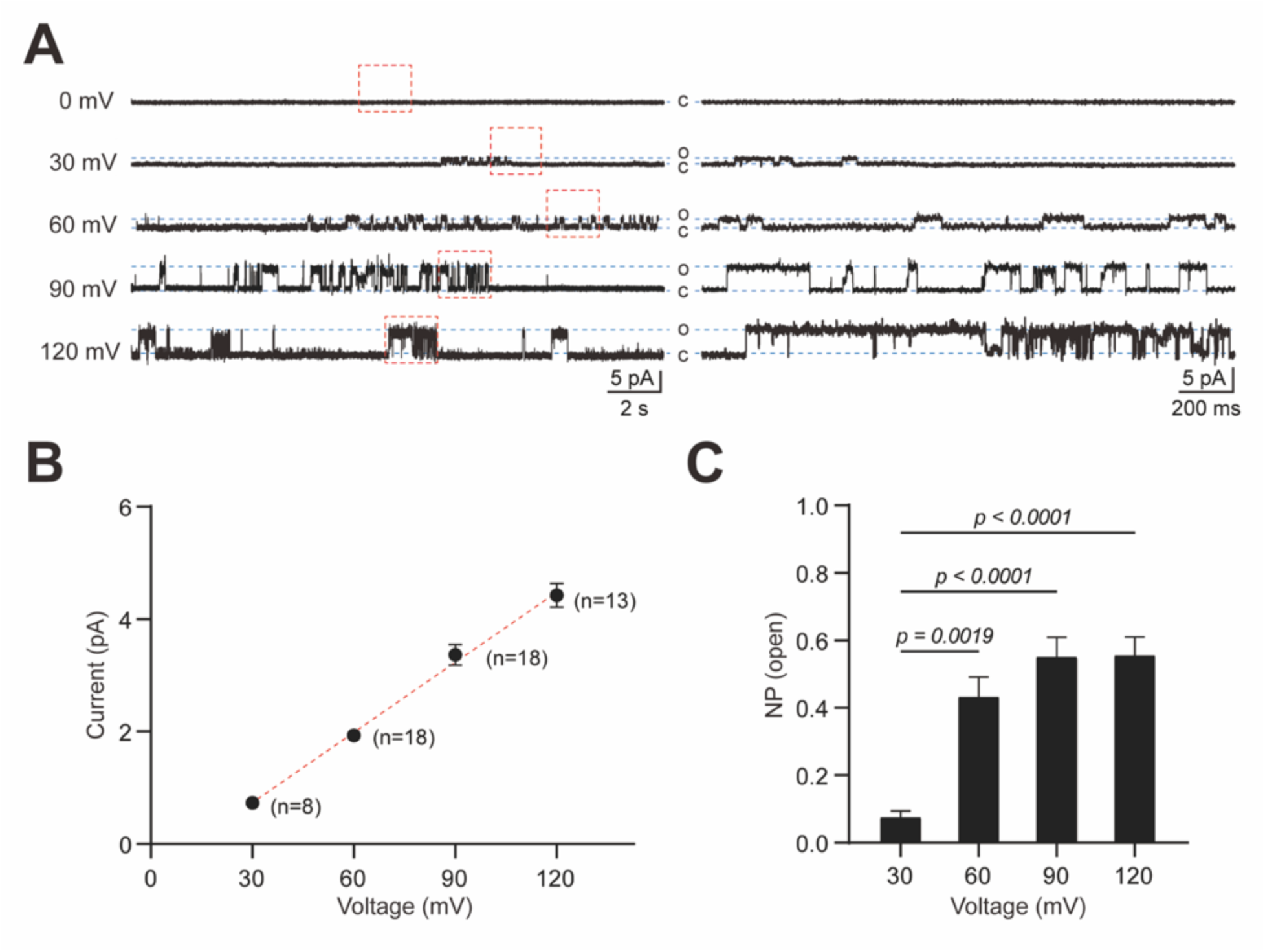
Single-channel recordings of VGAC channel. **(A)** Representative single-channel current traces in outside-out patch-clamp recordings from CHO-K1 cells expressing VGAC at 0 mV, 30 mV, 60 mV, 90 mV, and 120 mV. The traces shown on the right are enlargements of the traces within the dotted box on the left. **(B)** The current amplitudes were plotted against voltage potentials ranging from 30 mV to 120 mV in 30 mV increments, revealing an estimated slope conductance of 41.98±2.71 pS. The coefficient of determination (R^2^) is 0.8139. **(C)** The bars in the graph represent the nominal open probability (NPo) (mean ± SEM) recorded at 30 mV (n=8), 60 mV (n=18), 90 mV (n=18), and 120 mV (n=13). The p-values obtained from the one-way ANOVA (Analysis of Variance) followed by Tukey’s comparison test are reported.

Consistent with the voltage-dependent current observed in the whole-cell patch clamp experiments, gating in the single-channel recordings was visualized. At 30 mV, the nominal open probability of VGAC was very low (about 0.07). In sharp contrast, at 60 mV, the open probability of VGAC increased by approximately 6-fold to 0.43, and further increased to 0.55 at 90 mV (Figures 3A and 3C). At 120 mV, the open probability of VGAC plateaued and is similar to that of 90 mV. These results confirm that our designer VGAC channel is a voltage-gated ion channel. Voltage-gating is due to the dramatic increase in the open probability at high positive voltages, and the single-channel conductance remains constant.

### Cryo-EM structure of VGAC

Cryo-EM micrographs were collected for VGAC and processed following standard protocols (Figure S5). To mitigate potential bias, C1 symmetry was applied during automatic image processing and classification. From the analysis of 343,211 selected particles, we determined the cryo-EM structure of VGAC at a resolution of 2.9 Å (Figures 4A and S5, Table S2). The cryo-EM structure clearly demonstrates a pentameric pore assembly, consistent with the design model. Both the transmembrane domain and the cytoplasmic domain are nicely resolved (Figures 4A and S5). The linkers connecting the cytoplasmic domain and the transmembrane domain exhibit weak density in the EM map. The density surrounding the membrane-spanning region is likely derived from surrounding detergent molecules (Figure S5B).

**Figure 4.**
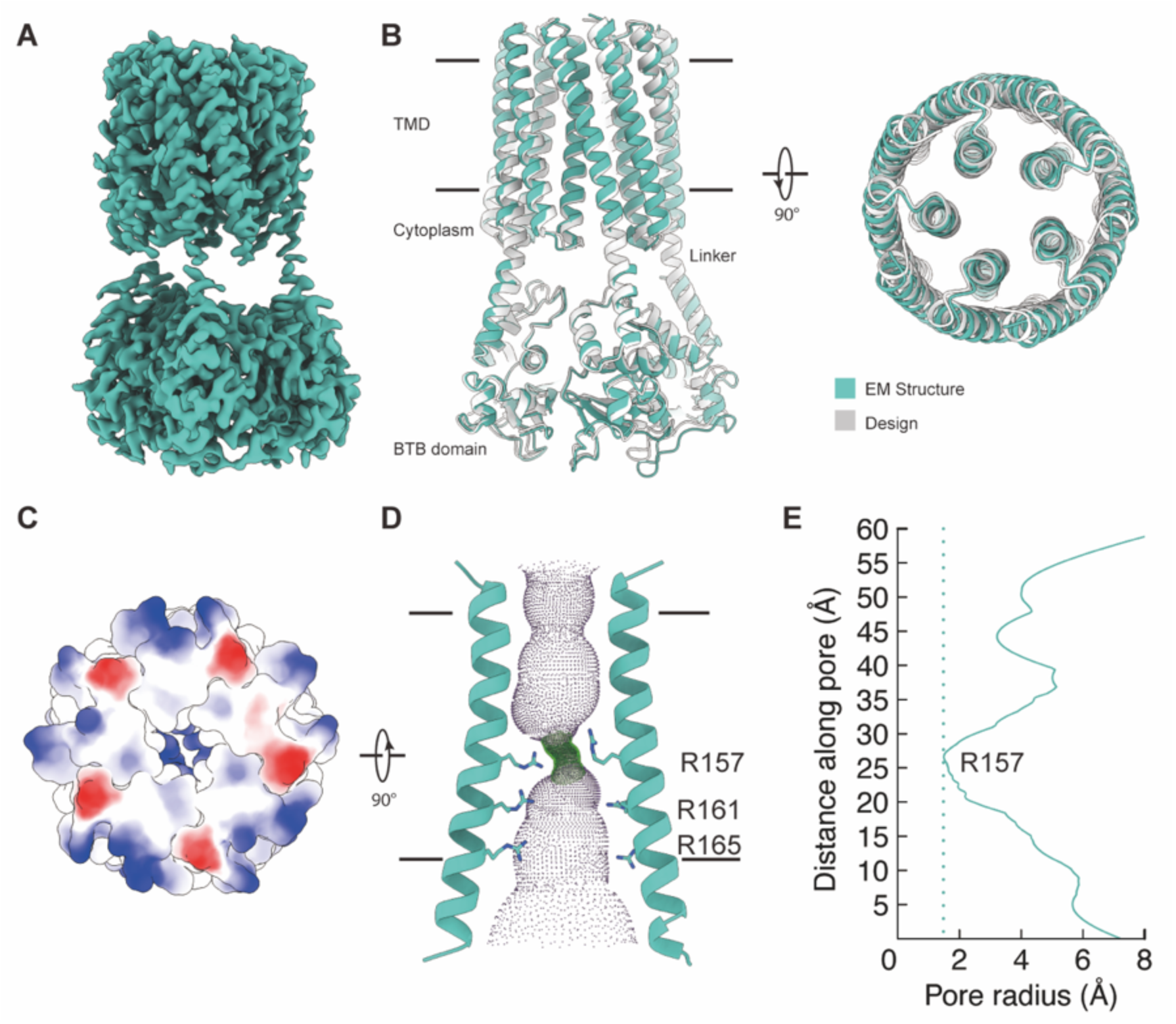
Cryo-EM structure of VGAC. **(A)** A cryo-EM density map of VGAC at a resolution at 2.9 Å. **(B)** Two perpendicular views of the superposition of the cryo-EM structure (shown in green) and the design model (shown in gray) of VGAC. TMD, transmembrane domain. **(C)** The electrostatic surface of VGAC, as viewed from the extracellular space, reveals a closed channel by the arginine constriction. **(D)** Pore profile of VGAC. Purple dots represent internal surface of VGAC channel, with side chains shown for R157, R161 and R165 from two of the five S2 α helices that line the pore. The helices are shown as a cartoon and the side chains are shown as sticks. **(E)** The pore radius along the central axis. Orientation and dimensions are as in **(D)**. See also Figure S2, S5 and Table S2.

The cryo-EM structure of the transmembrane domain of VGAC closely resembles the design model, with a Cα root-mean-square deviation (RMSD) of 1.09 Å for all aligned residues (Figure 4B). The 15-helix transmembrane domain exhibits a novel structure, as no similar structure was identified in a Dali search^36^ of the protein data bank (PDB) for the assembly.

As designed, the S2 helices form a funnel-like inner ring, expanding towards the cytoplasmic side and narrowing to form a short neck on the opposite side (Figures 4D and 4E). This configuration gives rise to a continuous pore that spans approximately 55 Å in length. Adjacent to the pore’s opening on the cytoplasmic side, there are five lateral openings situated between the BTB domain, transmembrane domain, and linkers. These lateral openings possess sufficient size to permit the passage of ions (Figures 4A and 4B).

The cryo-EM structure of VGAC revealed a central channel with a positively charged constriction that may attract anions (Figures 4C, 4D and S5G-S5L). The designed R157 is located approximately halfway in the transmembrane domain and forms the narrowest constriction in the pore with a radius of 1.49 Å, which is smaller than that of chloride ions (radius of ∼1.8 Å). Other parts along the central pore exhibit a wide-open pore with radii over 3 Å (Figure 4E). This data is consistent with whole-cell patch clamp experiments and suggests that the structure of VGAC represents a non-conducting closed state at the voltage of 0 mV, corresponding to the conditions under which the cryo-EM structures were determined (Figure 4C). The finding that the iodide ion was barely permeable through the pore indicates that the upper limit of the radius of the pore dilation is equal to the radius of the iodide ion (∼2.15 Å). This means that the dynamic range of pore dilation is 1.3 Å, from a diameter of 3.0 Å in the closed state to 4.3 Å in the fully open state that hardly passes iodide.

### Function of the arginine constriction in VGAC

We conducted single point mutations on each of the three Arg constriction residues in VGAC, to replace them with their corresponding residues in tmZC8-BTB. The R157G mutant exhibited a significantly decreased current compared to VGAC, registering a current of 405±21 pA at +120 mV (light blue line, Figure 5; Figure S4L; Table S1). The R161A mutant demonstrated current at positive potentials above 40 mV but also at negative potentials below −120 mV (-320±37 pA at - 120 mV) (pink line, Figure 5; Figure S4L; Table S1). This may indicate that removing a positive charge at this position resulted in a channel that lost its unidirectional ion transport and outward rectification. The R165L mutation displayed an I-V curve similar to that of VGAC (deep blue line, Figure 5; Figure S4L; Table S1). This indicates that R165 residue plays the least role in maintaining anion conductivity. Consequently, these findings indicate that R157 plays a critical role in controlling voltage gating and ion permeation, while R161 contributes to the unidirectional flow of anions by supporting the structure of the R157 constriction site.

**Figure 5.**
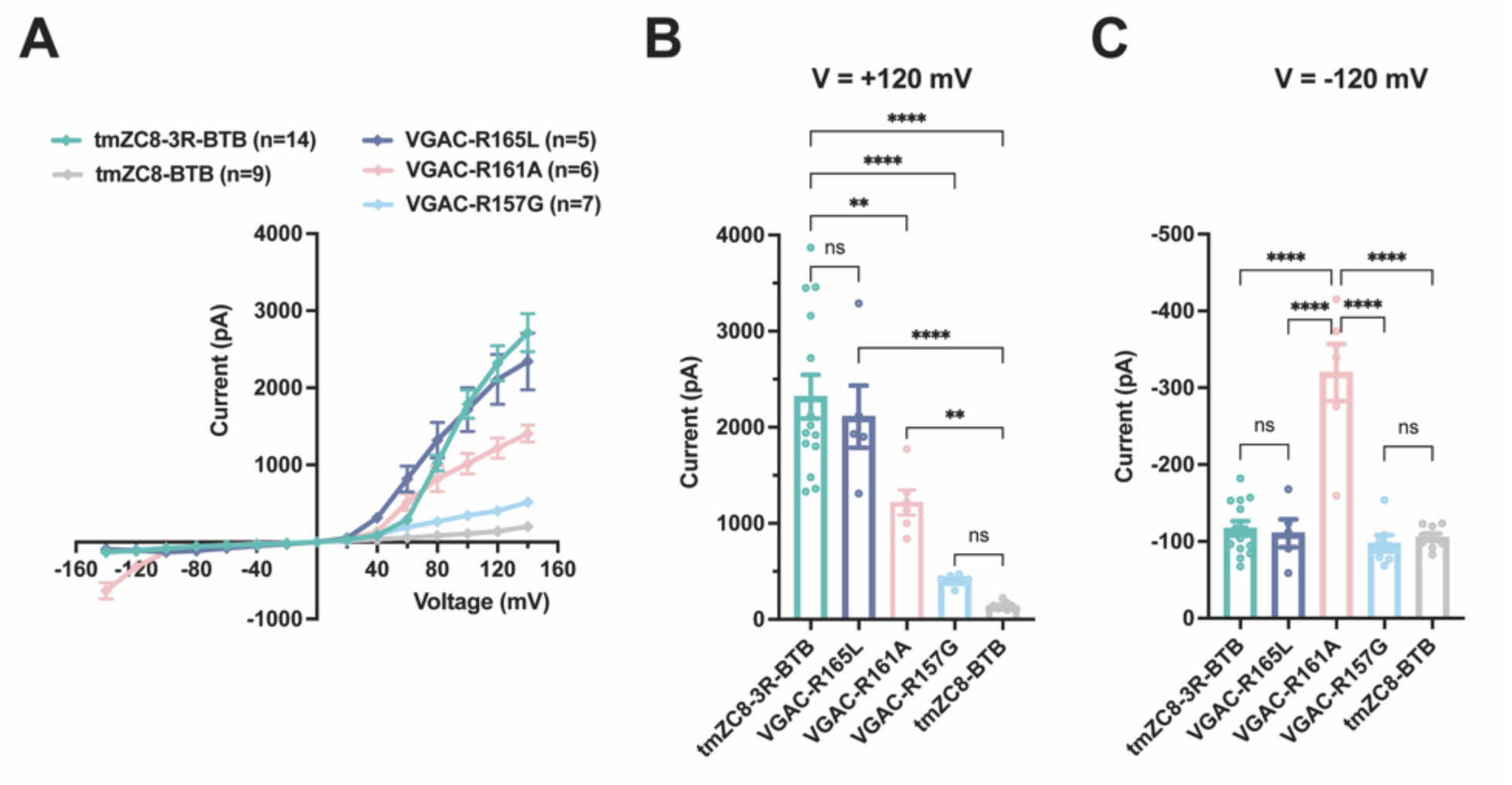
Function of the arginine constriction of VGAC. Conductivity of VGAC mutation variants. The whole-cell currents of cells transfected with VGAC and tmZC8-BTB are shown as controls. **(A)** I-V curves. **(B and C)** the bars represent the mean currents at 120 mV (B) and -120 mV (**C**), and the error bars depict the mean ± standard error of the mean (SEM). The P-values obtained from the one-way ANOVA (Analysis of Variance) followed by Tukey’s comparison test are reported (*p < 0.05, **p <= 0.01, ***p <= 0.001, ****p <= 0.0001, ns = no significance). See also Figure S4 and Table S1.

### Cryo-EM structure of tmZC8-BTB

It is puzzling that tmZC8-BTB did not show any current and the R157G mutant of VGAC exhibited a small current in the electrophysiology experiments, as the arginine constrictions in VGAC were replaced by small hydrophobic residues in tmZC8-BTB, likely resulting in a more open pore. To address this question, we solved the cryo-EM structure of tmZC8-BTB at a resolution of 3.3 Å, and the cryo-EM structure of the transmembrane domain of tmZC8-BTB closely resembles the design model, with a Cα root-mean-square deviation (RMSD) of 1.33 Å for all aligned residues (Figures 6A and S6, Table S2). The backbone structure of tmZC8-BTB also closely resembles that of VGAC, with an overall RMSD of 1.0 Å (Figure 6B). The transmembrane domain is composed of five nearly identical protomers, with only minor differences observed (Figure S7A). Notably, both within each tmZC8-BTB protomer and at the interface between individual subunits, interactions among core residues are predominantly hydrophobic packing. Side-chain hydrogen bonding and classic GXXXG motifs are absent in the core, which play crucial roles in membrane protein folding^37,38^ (Figures S7B and S7C).

**Figure 6.**
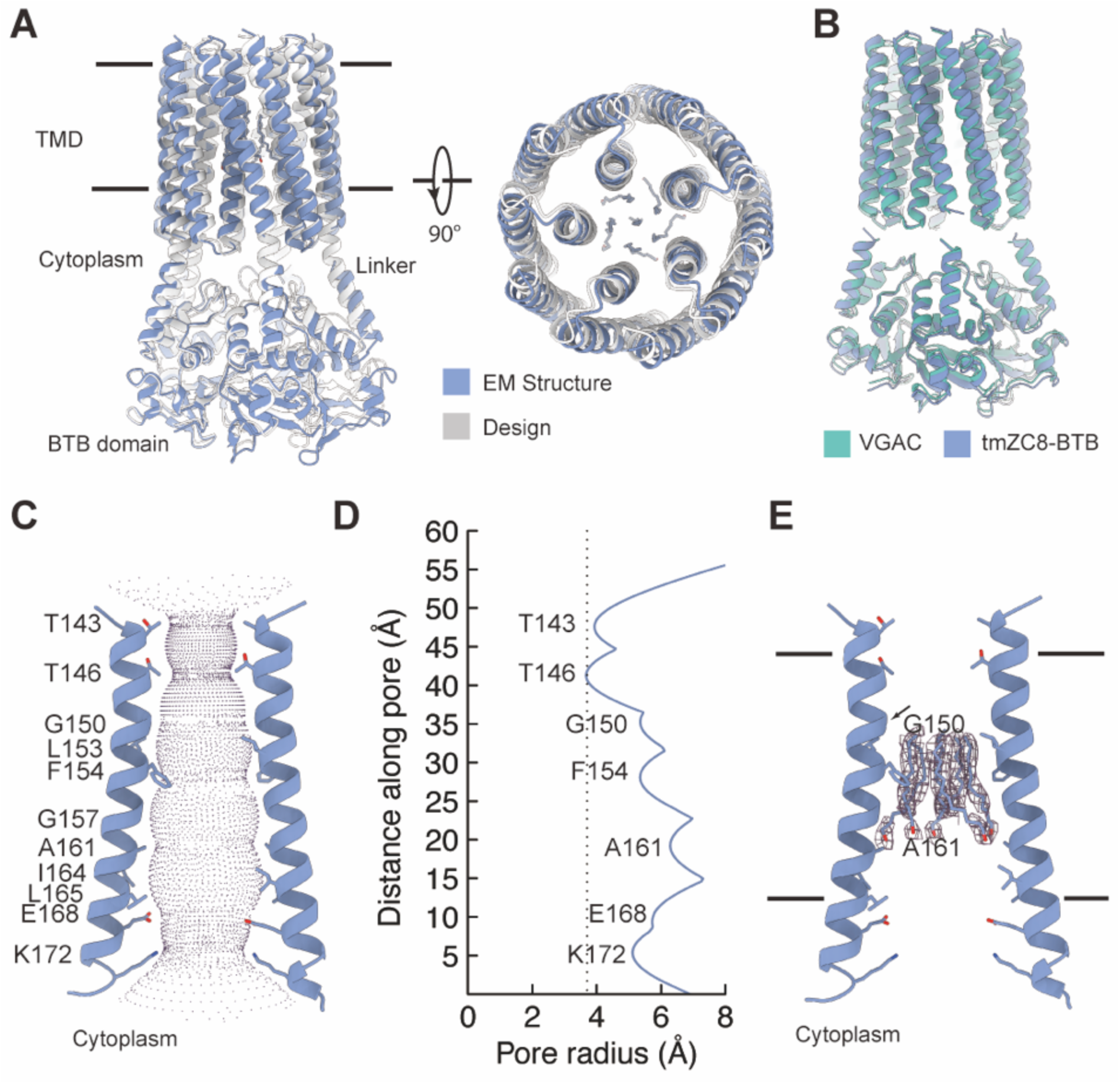
Cryo-EM structure of tmZC8-BTB. **(A)** Two views of the backbone superposition of tmZC8-BTB cryo-EM structure (blue) and design model (gray). TMD, transmembrane domain. The structure includes a bundle of six hydrophobic tails of DDM, represented as sticks. **(B)** Backbone superposition of cryo-EM structures of VGAC (green) and tmZC8-BTB (blue). **(C)** Pore profile of tmZC8-BTB. The pore of tmZC8-BTB is displayed in relation to two opposite subunits with pore-lining residues shown. The S2 helices are shown as a cartoon and the side chains of the pore-lining residues are shown as sticks. The BTB domain is removed for clarity. **(D)** The pore radius along the central axis. Orientation and dimensions are as in (**C**). **(E)** Detergent positions observed in the tmZC8-BTB structure. Detergents are shown as sticks with the cryo-EM map carved around. The position of the Cα atom of G150 is indicated by a black arrow. Orientation and dimensions are as in (**C**). See also Figure S2, S6, S7 and Table S2.

The cryo-EM structure of tmZC8-BTB clearly shows a channel structure. The pore-lining residues comprise four charged residues (E168, R169, K172 and E173) near the cytoplasmic opening of the pore, and two threonine residues (T143 and Thr146) on the opposite side (Figures 6C and 6D). T143 and T146 form the narrowest regions of the pore with radii of 4 and 3.7 Å, respectively. The pore-lining residues between T146 and E168 comprise three small amino acids (G150, G157, and A161), resulting in pore radii exceeding 5 Å in this region.

Interestingly, the cryo-EM map reveals a bundle of six tubular densities blocking the channel, among which the central tubular density is not as well-defined as the five peripheral ones (Figures 6A and 6E). These densities were modeled as the acyl chain tails of dodecyl-maltoside (DDM) molecules, considering their shape and chemical environment. The six acyl chains are positioned between the levels of G150 and A161, making extensive hydrophobic interactions with the internal surface of the pore. However, the density for the head groups of these DDM molecules is poorly resolved. The orientation of the polar heads of the five peripheral detergents was assigned to point towards the cytoplasmic side considering the space in the pore.

By comparing the structures of VGAC and tmZC8-BTB (Figures 4D and 6E), two major differences become apparent. Firstly, the presence of three Args in VGAC alters the geometry and electrostatic potential of the central channel. Secondly, in tmZC8-BTB, there is a detergent bundle that blocks the pore, whereas in the VGAC structure, this space is predominantly occupied by R157. One possible explanation for the non-conductivity of tmZC8-BTB and small conductivity of VGAC R157G mutant is that certain lipid molecules from the membrane may obstruct the hydrophobic channel, as observed in the cryo-EM structure of tmZC8-BTB with the DDM detergent.

## DISCUSSION

We have successfully *de novo* designed VGAC, a voltage-gated anion channel that undergoes custom-designed conformational changes induced by membrane potential. To our knowledge, VGAC represents the first example of *de novo* transmembrane proteins that can undergo conformational changes in response to a physical stimulus. Also, VGAC is the first accurately *de novo* designed ion channel confirmed by high-resolution three-dimensional structure in experiments, and the cryo-EM structure perfectly matches with the designed model. VGAC possesses a two-ring pentameric structure featuring a positively charged arginine constriction that serves as the voltage sensor, the selectivity filter, and the unidirectional valve for anions, setting it apart from native channels and yielding fresh insights into the mechanisms of voltage-gating and ion selectivity.

Our cryo-EM structure and electrophysiology data support a model in which the sidechains of R157 form a constriction that blocks ion conduction at 0 mV (Figure 7A). Increasing the potential above 40 mV causes an upward movement of the R157 sidechains, thereby opening an ion pathway through the channel (Figure 7B). Anions are attracted by the positive electrostatic environment in the pore and flow through the channel from the extracellular side. As I^−^ (radius of ∼2.15 Å) is larger than Cl^−^ (radius of ∼1.81 Å), the high selectivity for Cl^−^ ions over iodide ions (I^−^) suggests a narrow dynamic range of constriction dilation in VGAC and the arginine constriction may serve as selectivity filter based on anion sizes. We propose that further upward movement of R157 and greater pore dilation at higher membrane potential may be restricted by the funnel-shaped inner-ring backbone, which tapers upward. At negative potentials, the downward movement of R157 for opening VGAC channel is likely hampered by charge-charge repulsion and steric hindrance among R157 residues and the rest of the channel. Notably, R161 plays an important role in maintaining the unidirectional flow of anions, as elucidated by electrophysiology data from the R161A mutant.

**Figure 7.**
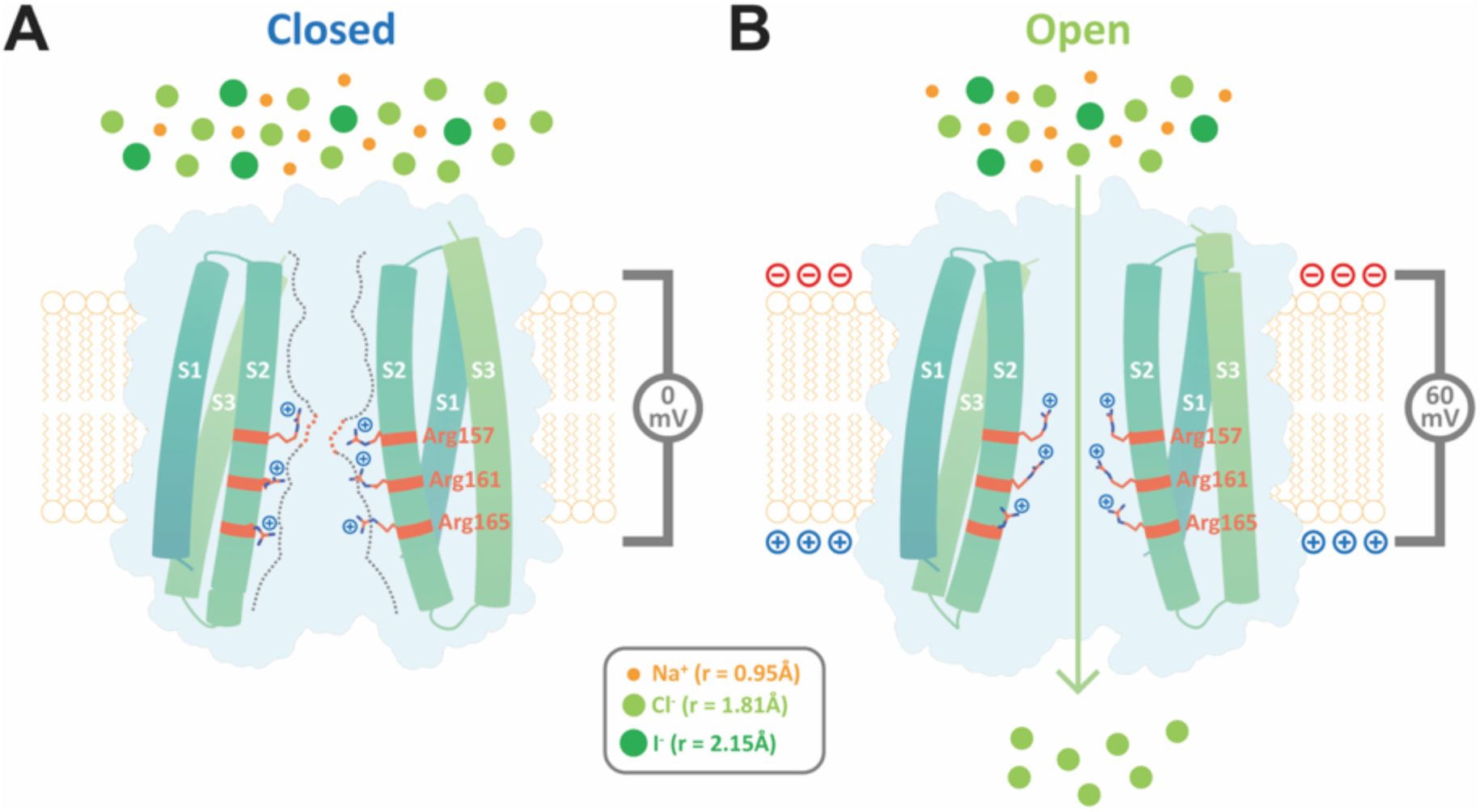
Model for voltage gating and ion selectivity of VGAC. **(A and B)** Model using two opposing subunits of VGAC. In a closed state (**A**), R157 form a seal to ion conduction, which is captured in the cryo-EM structure. Channel opening at membrane potentials above 40 mV (**B**) involves an upward movement of the side chains of R157. This movement causes the channel aperture to dilate, allowing for the passage of ions. Anions, specifically attracted by the arginine filter located within the pore, flow through the channel from the extracellular side. VGAC exhibits a strong preference for Cl^−^ over I^−^, indicating that additional pore dilation is limited in the funnel-shaped structure at more positive potentials.

It is instructive to compare the structure and function of our *de novo* designed VGAC with those of natural chloride channels and to illustrate the potential utility of VGAC. Firstly, VGAC exhibits a unique structure and primary sequence as there are no similar ones exist in nature, and VGAC possesses much simpler structure than natural ion channels. Secondly, the ion selectivity sequence of VGAC is distinct from that of most chloride channels^33,39–46^, which typically follow the Hofmeister lyotropic selectivity sequence of I^−^ > Br^−^ > Cl^−47^, where selectivity is largely determined by the energetic cost of anion dehydration. Although the ion selectivity sequence in VGAC is reminiscent of the voltage-gated ClC channels (Cl^−^ > Br^−^ >> I^−^), CLC channels structurally resemble Cl^−^/H^+^ transporters and the backbone amides are determinants of Cl^−^ selectivity^21,48,49^; while in VGAC, the arginine constriction selects for Cl^−^. Thirdly, the single-channel conductance of VGAC was measured to be 41.98±2.71 pS, which is considerably larger than the 1–1.5 pS of the ClC-1 channel ^50,51^. Interestingly, VGAC has a similar open-channel conductance to some ligand-gated chloride channels, such as the glycine receptor and GABAA

receptor, which range approximately from 35 to 50 pS^22^. Lastly, VGAC possesses a unique gating mechanism. Most of the chloride channels are gated by ligands, proton, calcium or hypotonic stress ^33,39–46^. For the voltage-gated CLC channel, voltage drives a permeant anion or a proton into the pore, thereby opening the glutamate gate through repulsion or protonation^52–54^. CLC-1 predominately exists in the open state at 0 mV and tends to close in response to negative transmembrane voltages, whereas CLC-2 opens in response to hyperpolarization and tends to close at 0 mV^55^. However, in VGAC, the arginine constriction likely undergoes voltage induced conformational changes for opening at potentials above 40 mV which allows the unidirectional influx of Cl^−^. Taken together, compared to natural anion channels, VGAC adopts a much simpler and more stable structure, and possesses distinct ion selectivity and gating mechanism. Due to the unique features of VGAC, it may serve as a molecular tool for reducing abnormal hyperactive neuronal activities in seizures and epilepsy, or for regulating the membrane potential of neutrophils. Neutrophils depolarize to +58 mV during the respiratory burst^56–58^, which can in turn regulate the production of superoxide. Also, VGAC may enable further design for tunable function. For example, the activation potential of VGAC may be further tuned down to the range of 0-20 mV to potentially regulate cardiac or neuronal activities.

VGAC incorporates novel features for *de novo* transmembrane proteins, including conformation changes, gating charges, a three-helix protomer, and a pentameric funnel-shaped helical bundle. It is quite surprising that a *de novo* transmembrane channel carries 15 positively charged arginine residues within the central pore without any counterbalance charges but still maintained a stable structure. In addition, VGAC adopts an Nin-C_out_ topology, which is distinct from the predominant N_in_-C_in_ topologies observed in both natural transmembrane proteins^59^ and previously designed multi-pass transmembrane proteins^2,6^. This topology allows for further protein engineering, such as the fusion of ligand-binding domains on either side of the membrane for the design of channel-based sensors. Moreover, the funnel-shaped helical bundle, assembled from three-helix protomers, represents a previously unexplored structural space in both *de novo* protein design and in nature.

We utilized deep learning-based methods^26,27,60^ to facilitate the design of core-residue packing and predict the structural integrity. Our results demonstrate that the ProteinMPNN^26^ deep network, primarily trained on water-soluble proteins, can effectively design core packings for transmembrane proteins. Notably, the hydrophobic core packing provides sufficient stabilizing interactions for the functional VGAC, even though the pore contains 15 Args. Our results suggest that *de novo* transmembrane proteins with a molecular weight of approximately 55 kDa, and possessing 15 transmembrane helices, can be designed through the packing of hydrophobic side chains. In addition, our design approach is more straightforward compared to previous methods, by reducing the laborious efforts in determining the structure of a water-soluble precursor of the transmembrane design^2,6,8^.

The ability to design ion channels with customized gating properties in response to specific environmental stimuli not only enriches our understanding of the fundamental principles of membrane biophysics but also unlocks a wide array of potential applications. VGAC is genetically encodable and functions within mammalian cells, thus laying the foundation for the creation of protein tools that capable of regulating membrane potentials in animals. In future studies, a wide variety of designer transmembrane proteins with tailored stimuli-induced conformational changes could conceivably play a role in the detection of specific ligands or physical stimuli both *in vitro* and *in vivo*.

## SUPPLEMENTAL INFORMATION

Document S1. Table S1 and Table S2

## ACKNOWLEDGEMENTS

We would like to thank the cryo-EM facility, microscopy facility, the HPC Center for computation assistance, and the Protein Characterization and Crystallography Facility of Westlake University for help. This work was funded by Ministry of Science and Technology of the People’s Republic of China project 2020YFA0909200 (P.L.), Zhejiang Provincial Natural Science Foundation of China Grant No. LR23C050001 (P.L.), “Pioneer” and “Leading Goose” R&D Program of Zhejiang (Grant no. 2024SSYS0036), National Natural Science Foundation of China project 22137005 (P.L.), and the Research Center for Industries of the Future (RCIF) at Westlake University.

## AUTHOR CONTRIBUTIONS

P.L. conceived the research; C.Z., H.L., J.W., K.W. and P.L. designed experiments; C.Z., H.L., and J.W. contributed equally to this work; C.Z. developed the computational methods and designed the proteins. H.L. conducted the biochemical experiments and collected cryo-EM data. H.L., C.Z. and S.S. conducted cloning experiments. C.Z. and K.S. solved the EM structures. Patch clamp experiments were performed by J.W. and Z.C., with guidance from K.W. A.Z., J.W., X.J., F.Y., and T.M.G. contributed to the patch clamp experiments. P.L. wrote the original draft and all authors participated in manuscript revision.

## REFERENCES

1. Joh, N.H., Wang, T., Bhate, M.P., Acharya, R., Wu, Y., Grabe, M., Hong, M., Grigoryan, G., and DeGrado, W.F. (2014). De novo design of a transmembrane Zn(2)(+)-transporting four-helix bundle. Science 346, 1520–1524. 10.1126/science.1261172.

2. Lu, P., Min, D., DiMaio, F., Wei, K.Y., Vahey, M.D., Boyken, S.E., Chen, Z., Fallas, J.A., Ueda, G., Sheffler, W., et al. (2018). Accurate computational design of multipass transmembrane proteins. Science 359, 1042–1046. 10.1126/science.aaq1739.

3. Mravic, M., Thomaston, J.L., Tucker, M., Solomon, P.E., Liu, L., and DeGrado, W.F. (2019). Packing of apolar side chains enables accurate design of highly stable membrane proteins. Science 363, 1418–1423. 10.1126/science.aav7541.

4. Vorobieva, A.A., White, P., Liang, B., Horne, J.E., Bera, A.K., Chow, C.M., Gerben, S., Marx, S., Kang, A., Stiving, A.Q., et al. (2021). De novo design of transmembrane β barrels. Science 371, eabc8182. 10.1126/science.abc8182.

5. Lear, J.D., Wasserman, Z.R., and DeGrado, W.F. (1988). Synthetic amphiphilic peptide models for protein ion channels. Science 240, 1177–1181. 10.1126/science.2453923.

6. Xu, C., Lu, P., Gamal El-Din, T.M., Pei, X.Y., Johnson, M.C., Uyeda, A., Bick, M.J., Xu, Q., Jiang, D., Bai, H., et al. (2020). Computational design of transmembrane pores. Nature 585, 129–134. 10.1038/s41586-020-2646-5.

7. Kratochvil, H.T., Watkins, L.C., Mravic, M., Thomaston, J.L., Nicoludis, J.M., Somberg, N.H., Liu, L., Hong, M., Voth, G.A., and DeGrado, W.F. (2023). Transient water wires mediate selective proton transport in designed channel proteins. Nat Chem 15, 1012–1021. 10.1038/s41557-023-01210-4.

8. Scott, A.J., Niitsu, A., Kratochvil, H.T., Lang, E.J.M., Sengel, J.T., Dawson, W.M., Mahendran, K.R., Mravic, M., Thomson, A.R., Brady, R.L., et al. (2021). Constructing ion channels from water-soluble alpha-helical barrels. Nat Chem 13, 643–650. 10.1038/s41557-021-00688-0.

9. Berhanu, S., Majumder, S., Müntener, T., Whitehouse, J., Berner, C., Bera, A.K., Kang, A., Liang, B., Khan, N., Sankaran, B., et al. (2024). Sculpting conducting nanopore size and shape through de novo protein design. Science 385, 282–288. 10.1126/science.adn3796.

10. Joh, N.H., Grigoryan, G., Wu, Y., and DeGrado, W.F. (2017). Design of self-assembling transmembrane helical bundles to elucidate principles required for membrane protein folding and ion transport. Philos Trans R Soc Lond B Biol Sci 372. 10.1098/rstb.2016.0214.

11. Niitsu, A., Heal, J.W., Fauland, K., Thomson, A.R., and Woolfson, D.N. (2017). Membrane-spanning alpha-helical barrels as tractable protein-design targets. Philos Trans R Soc Lond B Biol Sci 372. 10.1098/rstb.2016.0213.

12. Zhou, C., and Lu, P. (2022). De novo design of membrane transport proteins. Proteins 90, 1800–1806. 10.1002/prot.26336.

13. Zhu, J., and Lu, P. (2022). Computational design of transmembrane proteins. Curr Opin Struct Biol 74, 102381. 10.1016/j.sbi.2022.102381.

14. An, L., Said, M., Tran, L., Majumder, S., Goreshnik, I., Lee, G.R., Juergens, D., Dauparas, J., Anishchenko, I., Coventry, B., et al. (2024). Binding and sensing diverse small molecules using shape-complementary pseudocycles. Science 385, 276–282. 10.1126/science.adn3780.

15. Catterall, W.A. (1995). Structure and function of voltage-gated ion channels. Annu Rev Biochem 64, 493–531. 10.1146/annurev.bi.64.070195.002425.

16. Jiang, Y., Lee, A., Chen, J., Ruta, V., Cadene, M., Chait, B.T., and MacKinnon, R. (2003). X-ray structure of a voltage-dependent K+ channel. Nature 423, 33–41. 10.1038/nature01580.

17. Shen, H., Zhou, Q., Pan, X., Li, Z., Wu, J., and Yan, N. (2017). Structure of a eukaryotic voltage-gated sodium channel at near-atomic resolution. Science 355. 10.1126/science.aal4326.

18. Yao, X., Gao, S., and Yan, N. (2024). Structural biology of voltage-gated calcium channels. Channels (Austin) 18, 2290807. 10.1080/19336950.2023.2290807.

19. Sands, Z., Grottesi, A., and Sansom, M.S. (2005). Voltage-gated ion channels. Curr Biol 15, R44–47. 10.1016/j.cub.2004.12.050.

20. Jentsch, T.J., and Pusch, M. (2018). CLC Chloride Channels and Transporters: Structure, Function, Physiology, and Disease. Physiol Rev 98, 1493–1590. 10.1152/physrev.00047.2017.

21. Dutzler, R., Campbell, E.B., Cadene, M., Chait, B.T., and MacKinnon, R. (2002). X-ray structure of a ClC chloride channel at 3.0 A reveals the molecular basis of anion selectivity. Nature 415, 287–294. 10.1038/415287a.

22. Waxham, M.N. (2004). CHAPTER 11 - Neurotransmitter Receptors. In From Molecules to Networks, J.H. Byrne, and J.L. Roberts, eds. (Academic Press), pp. 299–334. 10.1016/B978-012148660-0/50012-3.

23. Grigoryan, G., and Degrado, W.F. (2011). Probing designability via a generalized model of helical bundle geometry. J Mol Biol 405, 1079–1100. 10.1016/j.jmb.2010.08.058.

24. Huang, P.S., Oberdorfer, G., Xu, C., Pei, X.Y., Nannenga, B.L., Rogers, J.M., DiMaio, F., Gonen, T., Luisi, B., and Baker, D. (2014). High thermodynamic stability of parametrically designed helical bundles. Science 346, 481–485. 10.1126/science.1257481.

25. Thomson, A.R., Wood, C.W., Burton, A.J., Bartlett, G.J., Sessions, R.B., Brady, R.L., and Woolfson, D.N. (2014). Computational design of water-soluble alpha-helical barrels. Science 346, 485–488. 10.1126/science.1257452.

26. Dauparas, J., Anishchenko, I., Bennett, N., Bai, H., Ragotte, R.J., Milles, L.F., Wicky, B.I.M., Courbet, A., de Haas, R.J., Bethel, N., et al. (2022). Robust deep learning-based protein sequence design using ProteinMPNN. Science 378, 49–56. 10.1126/science.add2187.

27. Jumper, J., Evans, R., Pritzel, A., Green, T., Figurnov, M., Ronneberger, O., Tunyasuvunakool, K., Bates, R., Zidek, A., Potapenko, A., et al. (2021). Highly accurate protein structure prediction with AlphaFold. Nature 596, 583–589. 10.1038/s41586-021-03819-2.

28. Huang, P.S., Ban, Y.E., Richter, F., Andre, I., Vernon, R., Schief, W.R., and Baker, D. (2011). RosettaRemodel: a generalized framework for flexible backbone protein design. PLoS One 6, e24109. 10.1371/journal.pone.0024109.

29. Altschul, S.F., Gish, W., Miller, W., Myers, E.W., and Lipman, D.J. (1990). Basic local alignment search tool. J Mol Biol 215, 403–410. 10.1016/S0022-2836(05)80360-2.

30. Ji, A.X., Chu, A., Nielsen, T.K., Benlekbir, S., Rubinstein, J.L., and Privé, G.G. (2016). Structural Insights into KCTD Protein Assembly and Cullin3 Recognition. J Mol Biol 428, 92–107. 10.1016/j.jmb.2015.08.019.

31. Moller, S., Croning, M.D., and Apweiler, R. (2001). Evaluation of methods for the prediction of membrane spanning regions. Bioinformatics 17, 646–653. 10.1093/bioinformatics/17.7.646.

32. Schewe, M., Nematian-Ardestani, E., Sun, H., Musinszki, M., Cordeiro, S., Bucci, G., de Groot, B.L., Tucker, S.J., Rapedius, M., and Baukrowitz, T. (2016). A Non-canonical Voltage-Sensing Mechanism Controls Gating in K2P K(+) Channels. Cell 164, 937–949. 10.1016/j.cell.2016.02.002.

33. Ruan, Z., Osei-Owusu, J., Du, J., Qiu, Z., and Lu, W. (2020). Structures and pH-sensing mechanism of the proton-activated chloride channel. Nature 588, 350–354. 10.1038/s41586-020-2875-7.

34. Wang, Z., Wong, N.C., Cheng, Y., Kehl, S.J., and Fedida, D. (2009). Control of voltage-gated K+ channel permeability to NMDG+ by a residue at the outer pore. J Gen Physiol 133, 361–374. 10.1085/jgp.200810139.

35. Halm, D.R. (1998). Identifying swelling-activated channels from ion selectivity patterns. J Gen Physiol 112, 369–371. 10.1085/jgp.112.3.369.

36. Holm, L., and Sander, C. (1995). Dali: a network tool for protein structure comparison. Trends Biochem Sci 20, 478–480. 10.1016/s0968-0004(00)89105-7.

37. Joh, N.H., Min, A., Faham, S., Whitelegge, J.P., Yang, D., Woods, V.L., and Bowie, J.U. (2008). Modest stabilization by most hydrogen-bonded side-chain interactions in membrane proteins. Nature 453, 1266–1270. 10.1038/nature06977.

38. Teese, M.G., and Langosch, D. (2015). Role of GxxxG Motifs in Transmembrane Domain Interactions. Biochemistry 54, 5125–5135. 10.1021/acs.biochem.5b00495.

39. Chen, Y.H., Hu, L., Punta, M., Bruni, R., Hillerich, B., Kloss, B., Rost, B., Love, J., Siegelbaum, S.A., and Hendrickson, W.A. (2010). Homologue structure of the SLAC1 anion channel for closing stomata in leaves. Nature 467, 1074–1080. 10.1038/nature09487.

40. Hibbs, R.E., and Gouaux, E. (2011). Principles of activation and permeation in an anion-selective Cys-loop receptor. Nature 474, 54–60. 10.1038/nature10139.

41. Kane Dickson, V., Pedi, L., and Long, S.B. (2014). Structure and insights into the function of a Ca(2+)-activated Cl(-) channel. Nature 516, 213–218. 10.1038/nature13913.

42. Dang, S., Feng, S., Tien, J., Peters, C.J., Bulkley, D., Lolicato, M., Zhao, J., Zuberbuhler, K., Ye, W., Qi, L., et al. (2017). Cryo-EM structures of the TMEM16A calcium-activated chloride channel. Nature 552, 426–429. 10.1038/nature25024.

43. Liu, F., Zhang, Z., Csanady, L., Gadsby, D.C., and Chen, J. (2017). Molecular Structure of the Human CFTR Ion Channel. Cell 169, 85–95 e88. 10.1016/j.cell.2017.02.024.

44. Deneka, D., Sawicka, M., Lam, A.K.M., Paulino, C., and Dutzler, R. (2018). Structure of a volume- regulated anion channel of the LRRC8 family. Nature 558, 254–259. 10.1038/s41586-018-0134-y.

45. Wang, C., Polovitskaya, M.M., Delgado, B.D., Jentsch, T.J., and Long, S.B. (2022). Gating choreography and mechanism of the human proton-activated chloride channel ASOR. Sci Adv 8, eabm3942. 10.1126/sciadv.abm3942.

46. Liu, H., Polovitskaya, M.M., Yang, L., Li, M., Li, H., Han, Z., Wu, J., Zhang, Q., Jentsch, T.J., and Liao, J. (2023). Structural insights into anion selectivity and activation mechanism of LRRC8 volume- regulated anion channels. Cell Rep 42, 112926. 10.1016/j.celrep.2023.112926.

47. Hille, B.v., ed. (2001). Ion Channels of Excitable Membranes, 3rd edn.

48. Leisle, L., Lam, K., Dehghani-Ghahnaviyeh, S., Fortea, E., Galpin, J.D., Ahern, C.A., Tajkhorshid, E., and Accardi, A. (2022). Backbone amides are determinants of Cl(-) selectivity in CLC ion channels. Nat Commun 13, 7508. 10.1038/s41467-022-35279-1.

49. Wang, K., Preisler, S.S., Zhang, L., Cui, Y., Missel, J.W., Grønberg, C., Gotfryd, K., Lindahl, E., Andersson, M., Calloe, K., et al. (2019). Structure of the human ClC-1 chloride channel. PLOS Biology 17, e3000218. 10.1371/journal.pbio.3000218.

50. Pusch, M., Ludewig, U., Rehfeldt, A., and Jentsch, T.J. (1995). Gating of the voltage-dependent chloride channel CIC-0 by the permeant anion. Nature 373, 527–531. 10.1038/373527a0.

51. Pusch, M., Steinmeyer, K., and Jentsch, T.J. (1994). Low single channel conductance of the major skeletal muscle chloride channel, ClC-1. Biophys J 66, 149–152. 10.1016/S0006-3495(94)80753-2.

52. Pusch, M., Ludewig, U., Rehfeldt, A., and Jentsch, T.J. (1995). Gating of the voltage-dependent chloride channel CIC-0 by the permeant anion. Nature 373, 527–531. 10.1038/373527a0.

53. Dutzler, R., Campbell, E.B., and MacKinnon, R. (2003). Gating the Selectivity Filter in ClC Chloride Channels. Science 300, 108–112. 10.1126/science.1082708.

54. De Jesús-Pérez, J.J., Méndez-Maldonado, G.A., López-Romero, A.E., Esparza-Jasso, D., González- Hernández, I.L., De la Rosa, V., Gastélum-Garibaldi, R., Sánchez-Rodríguez, J.E., and Arreola, J. (2021). Electro-steric opening of the clc-2 chloride channel gate. Scientific Reports 11, 13127. 10.1038/s41598-021-92247-3.

55. Xu, M., Neelands, T., Powers, A.S., Liu, Y., Miller, S.D., Pintilie, G., Bois, J.D., Dror, R.O., Chiu, W., and Maduke, M. (2023). CryoEM structures of the human CLC-2 voltage gated chloride channel reveal a ball and chain gating mechanism. Cold Spring Harbor Laboratory.

56. DeCoursey, T.E., Morgan, D., and Cherny, V.V. (2003). The voltage dependence of NADPH oxidase reveals why phagocytes need proton channels. Nature 422, 531–534. 10.1038/nature01523.

57. Sasaki, M., Takagi, M., and Okamura, Y. (2006). A Voltage Sensor-Domain Protein Is a Voltage- Gated Proton Channel. Science 312, 589–592. 10.1126/science.1122352.

58. Ramsey, I.S., Moran, M.M., Chong, J.A., and Clapham, D.E. (2006). A voltage-gated proton- selective channel lacking the pore domain. Nature 440, 1213–1216. 10.1038/nature04700.

59. Daley, D.O., Rapp, M., Granseth, E., Melén, K., Drew, D., and von Heijne, G. (2005). Global topology analysis of the Escherichia coli inner membrane proteome. Science 308, 1321–1323. 10.1126/science.1109730.

60. Baek, M., DiMaio, F., Anishchenko, I., Dauparas, J., Ovchinnikov, S., Lee, G.R., Wang, J., Cong, Q., Kinch, L.N., Schaeffer, R.D., et al. (2021). Accurate prediction of protein structures and interactions using a three-track neural network. Science 373, 871–876. 10.1126/science.abj8754.

